# Mechanistic insights into Phage SSB-activated bacterial Retron-Eco8 immunity

**DOI:** 10.1101/2025.11.11.687766

**Authors:** Chao-Guang Ji, Zhuolin Li, Xin-Yang Wei, Yuelong Li, Jun-Tao Zhang, Xueyan Liu, Ning Jia

## Abstract

The Retron-Eco8 system, comprising a reverse transcriptase (RT), a non-coding RNA (ncRNA), and an OLD-family nuclease effector, protects bacteria from phage infection via abortive infection upon sensing a phage single-stranded DNA-binding protein (SSB). However, the molecular basis of this immunity remained unclear. Here, we report cryo-electron microscopy (cryo-EM) structures of Retron-Eco8 in inactive and activated states, revealing mechanisms of phage-triggered activation and effector function. Retron-Eco8 assembles into a tetrameric complex in which each protomer contains an RT, msrRNA–msdDNA duplex, and effector in an autoinhibited conformation. Upon phage infection, phage SSB binds msdDNA, relieving autoinhibition and activating the nuclease effector to degrade both phage and host DNA, triggering cell death to block phage propagation. Host SSB fails to activate the system, while DNA binding and oligomerization of phage SSB are essential for this activation, highlighting its specificity. These findings elucidate the molecular mechanism of Retron-Eco8-mediated immunity, facilitating retron-based biotechnological applications.

## Introduction

Reverse transcriptases (RTs) catalyze the synthesis of DNA from RNA templates. Retrons, typically comprising a reverse transcriptase (RT), a non-coding RNA (ncRNA), and an effector protein, represent a diverse class of multigene antiphage defense systems^1,2^. The ncRNA includes two inverted repeats (IRa1 and IRa2) that flank the internal *msr* and *msd* regions. The *msd* RNA serves as a template for reverse transcription by the retron RT, generating the DNA portion (msdDNA) that remains covalently linked to a conserved guanosine residue within the *msr* region through a 2′ –5′ phosphodiester bond. Reverse transcription terminates at a defined site, yielding a chimeric RNA-DNA heteroduplex that is subsequently processed by RNase H to form the mature msDNA-RNA duplex, known as multicopy single-stranded DNA (msDNA) (Fig. 1a)^2–4^. The ability of retrons to produce intracellular DNA molecules from structured ncRNAs makes them attractive tools for programmable genome engineering^5–7^. When coupled with single-stranded annealing proteins (SSAPs) and single-stranded binding proteins (SSBs), engineered retrons—termed recombitrons— enable efficient, markerless genome modification across multiple bacteriophages without counterselection^8^. Beyond genome editing, retrons have also been exploited as DNA-based recorders that can record cellular events^9,10^.

**Fig. 1.**
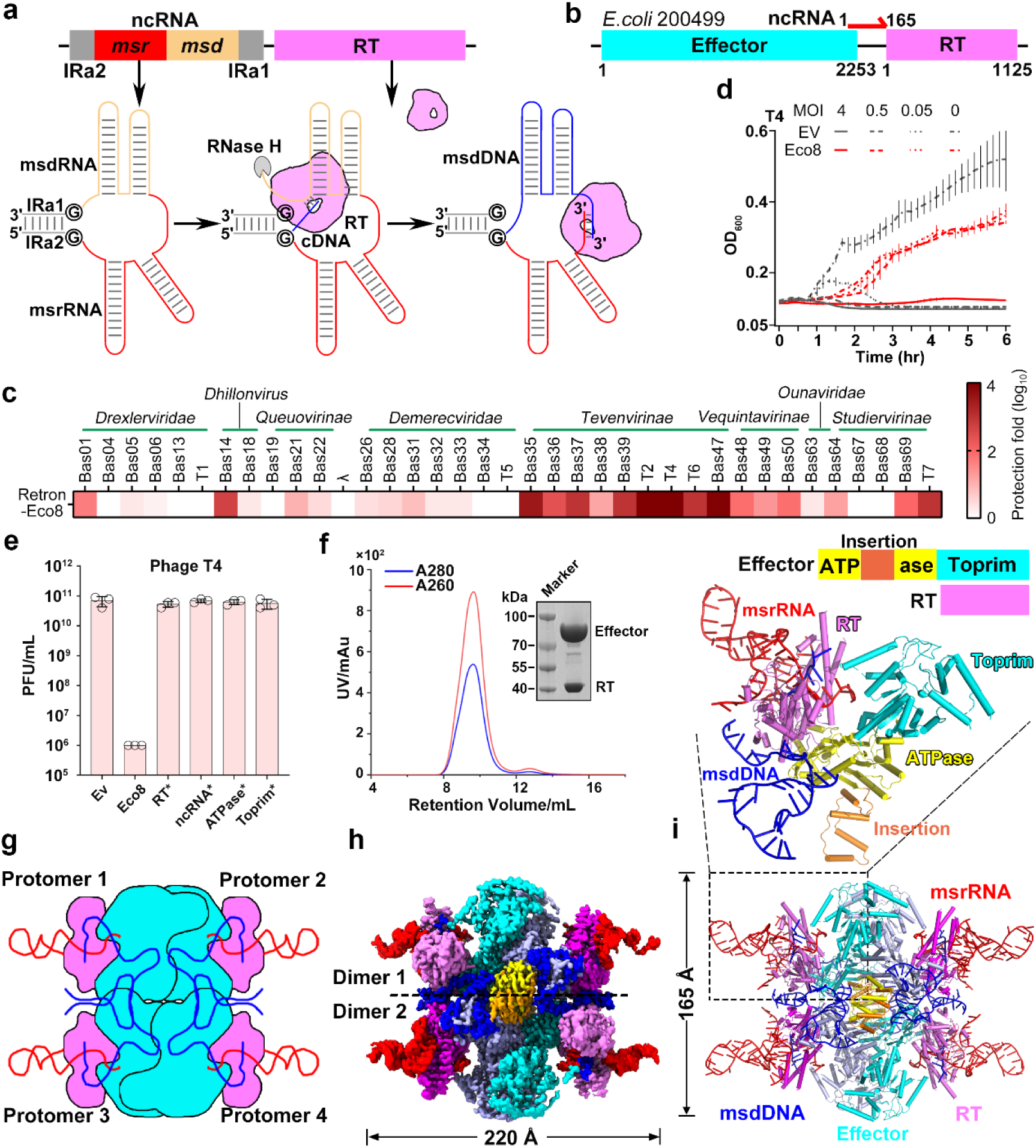
Overall architecture of the Retron-Eco8 tetramer. **a** Schematic depiction of the Retron reverse transcription process. **b** Schematic diagram of the genetic locus encoding effector, ncRNA and the reverse transcriptase found in *E. coli* strain 200499. **c** Plaque assays showing the resistance of *E. coli* strain MG1655 transformed with the Retron-Eco8 system or an empty pACYC vector (EV) against a wide range of phages. Images are representative of three biological replicates. **d** Growth curves of *E. coli* strain MG1655 in liquid culture transformed with the pACYC-Eco8 system or an empty pACYC vector (EV) infected by T4 phages. Bacteria were infected at time 0 at an MOI of 0 (no phage), 0.05, 0.5, or 4. Data are representative of three biological replicates. (n = 3; mean ± SD). **e** Mutational analysis of elements within the Retron-Eco8 defense system. Shown are plaque assays with phage T4, comparing the strain with wild-type system to with (RT*: YADD; ncRNA*: branching G; ATPase*: Walker B; Toprim*: DxD) mutants. Data are representative of three biological replicates. (n = 3; mean ± SD). **f** SEC and SDS-PAGE profiles show that the Retron-Eco8 complex assembles into a stable complex. **g-i** Schematic (g), Surface (h) and ribbon(i) representations of the 2.81-Å cryo-EM structure of the Retron-Eco8^ATP^ complex.

As antiphage defense systems, retrons safeguard bacterial hosts through diverse effector proteins, such as metabolic enzymes, membrane proteins, and nucleases^2^. In these systems, the RT together with msDNA sequester the toxic effector in an inactive state until phage infection triggers its activation, leading to abortive infection and host cell death. Elucidating the molecular diversity and activation mechanisms of retrons is therefore essential for both understanding bacterial immunity and harnessing these systems in biotechnology. The Retron-Eco8 system consists of a predicted OLD-family endonuclease effector, an RT, and a large intergenic ncRNA region that collectively mediate abortive infection upon phage invasion. Recent work revealed that the phage single-stranded DNA-binding protein (SSB) interacts with Retron-Eco8 and activates the system, triggering abortive infection to prevent phage propagation *in vivo*^2,3^. However, the molecular basis underlying system activation and how recognition of phage SSB triggers abortive infection remains unclear.

In this study, we used cryo-EM together with *in vitro* and *in vivo* analyses to elucidate the molecular architecture, activation, and effector mechanisms of the Retron- Eco8 complex. The RT and effector assemble into a stable heterotetrameric complex encased by the msrRNA–msdDNA duplex. Our structural and biochemical analyses further demonstrate that T7 SSB binds specifically to msdDNA, triggering system activation and inducing nonspecific degradation of intracellular DNA, including both phage and host DNA, ultimately leading to programmed cell death. Both DNA binding and oligomerization of T7 SSB are indispensable for Retron-Eco8 activation, whereas host SSB fails to trigger the process, highlighting the specificity of phage-derived SSB. These findings uncover the molecular principles governing retron activation and effector function, deepening our understanding of bacterial antiphage immunity and facilitating future biotechnological applications.

## Results

### Retron-Eco8 oligomerizes to mediate anti-phage defense through programmed cell death

To investigate the molecular mechanism underlying anti-phage defense by the Retron- Eco8 system, we reconstituted all three components–the effector protein, reverse transcriptase (RT), and non-coding RNA (ncRNA)–in *Escherichia coli* strain MG1655 and challenged the cells with a diverse panel of phages from the BASEL collection (Fig. 1b, c)^11^. Retron-Eco8 conferred resistance against a broad spectrum of phages, particularly those in the *Tevenvirinae* family, with up to a ∼ 10^4^-fold reduction in efficiency of plating (EOP) (Fig. 1c). Phage resistance was observed only at low multiplicities of infections (MOI) (MOI=0.05, 0.5), but was compromised at high MOI (MOI=4) (Fig. 1d), consistent with an abortive infection mechanism as previously described^2^. To determine whether the infected cells could resume growth, we plated Retron-Eco8-expressing and non-expressing cells infected with T4 phages at varying MOIs (0, 0.5, 5) and monitored the colony-forming units (CFUs) at multiple time points after infection. Cells expressing Retron-Eco8 failed to recover at high MOI, indicating that activation of the system triggers irreversible cell death (Extended Data Fig. 1). Mutations in conserved motifs of each component—including the YADD motif (Y198– D201) of RT (YADD to YAAA), DxD (D450–D452; DxD to AxA) and Walker B motifs (D285, D286; DD to AA) of the effector, and the branching guanosine of the ncRNA (G18C)—abolished defense activity (Fig. 1e), confirming that all three elements are essential for activity, in line with previous observations^2^. We next sought to determine whether these components form a stable complex. Co-expression of RT, ncRNA, and effector followed by size-exclusion chromatography (SEC) revealed a high-molecular-weight fraction with an elevated A260/A280 ratio, indicating the presence of RNA and the formation of an oligomer (Fig. 1f). Together, these findings indicate that Retron-Eco8 assembles into an oligomeric ribonucleoprotein complex that mediates abortive infection via programmed cell death, thereby limiting phage propagation.

### Retron-Eco8 adopts a tetrameric architecture

To gain mechanistic insights into Retron-Eco8-mediated anti-phage defense, we performed cryo-EM studies on SEC-purified oligomeric complexes. Two constructs were examined: one containing an N-terminal His₆ tag on the effector and a C-terminal Strep II tag on the reverse transcriptase (RT), and another carrying only the C-terminal Strep II tag on the RT and supplemented with Mg²⁺ and ATP. These complexes were resolved to overall resolutions of 2.94 Å and 2.81 Å, respectively. Structural interpretation focused on the latter dataset, which exhibited more complete density (Fig. 1g–i and Extended Data Figs. 2, 3a, b). The overall structure adopts a tetrameric assembly with a distorted "I"-shaped architecture, measuring approximately 220 Å in length, 165 Å in width, and 65 Å in height (Fig. 1h). The tetramer is organized as a dimer of dimers, with four effector subunits forming a central scaffold, each flanked by a peripherally associated reverse transcriptase (Fig. 1g–i). The msrRNA wraps around the RT, with its 3′ end reentering the conserved catalytic pocket and base-pairing with the reverse-transcribed msdDNA (Fig. 1i, upper inset). Despite the inclusion of ATP in the sample, densities corresponding to the ATP are unobserved.

Structural homology searches using DALI revealed that the Retron-Eco8 effector shares close similarity with GajA (PDB: 8X5I; Z-score = 16.3; Cα root mean square deviation (RMSD) = 4.5 Å over 398 residues)^12^, a member of the Overcoming Lysogenization Defect (OLD) family of endonucleases (Extended Data Fig. 3c)¹⁵. Like GajA, the effector contains an N-terminal ATPase domain (residues 1–390) and a C-terminal topoisomerase–primase (Toprim) domain (residues 391–750). The ATPase domain is further divided into two parts by a central α-helices insertion (161–256) (Fig. 1i and Extended Data Fig. 3c). The RT adopts a canonical right-hand-like fold and shows close structural similarity to the RT of *E. coli* Retron-Eco1 (Ec86) (PDB: 7V9U)^4^, with a RMSD of 2.1 Å (Extended Data Fig. 3d). The RT can be subdivided into three domains: fingers (residues 1–100 and 115–167), palm (residues 101–114 and 168–269), and thumb (residues 270–374) domains (Fig. 2a, b).

**Fig. 2.**
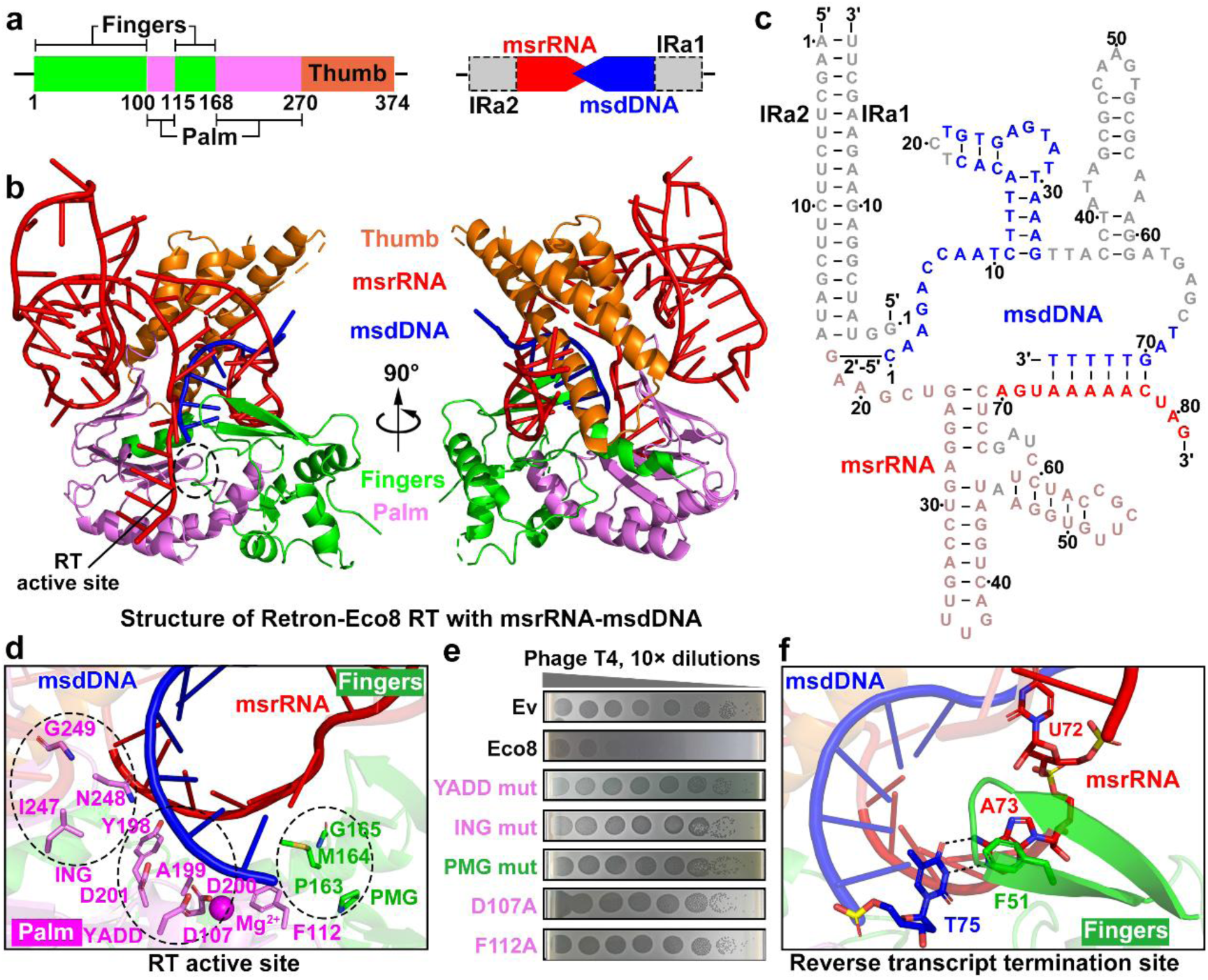
Structure of Retron-Eco8 RT protein and its associated msrRNA- msdDNA . **a, b** Domain organization (a) and ribbon representation (b) of the RT protein and msrRNA-msdDNA. **c** Secondary structure of Retron-Eco8 msrRNA-msdDNA. The regions not observed in the structure are colored in grey, and the msrRNA region predicted by AF3 and fitted into the map is colored in light red. **d** Close-up view of the RT active site in the Retron-Eco8 system. Residues are color-coded according to domain. **e** Serial dilution plaque assays for T4 phage on *E. coli* MG1655 strain transformed with plasmids encoding WT or indicated RT active site mutants. (YADD mut: 198YADD201/198AAAA201; ING mut:247ING249/247GGP249; PMG mut:163PMG165/163GPP165). **f** Detailed interactions of the reverse transcript termination site. Residues are color-coded according to domain.

**Fig. 3.**
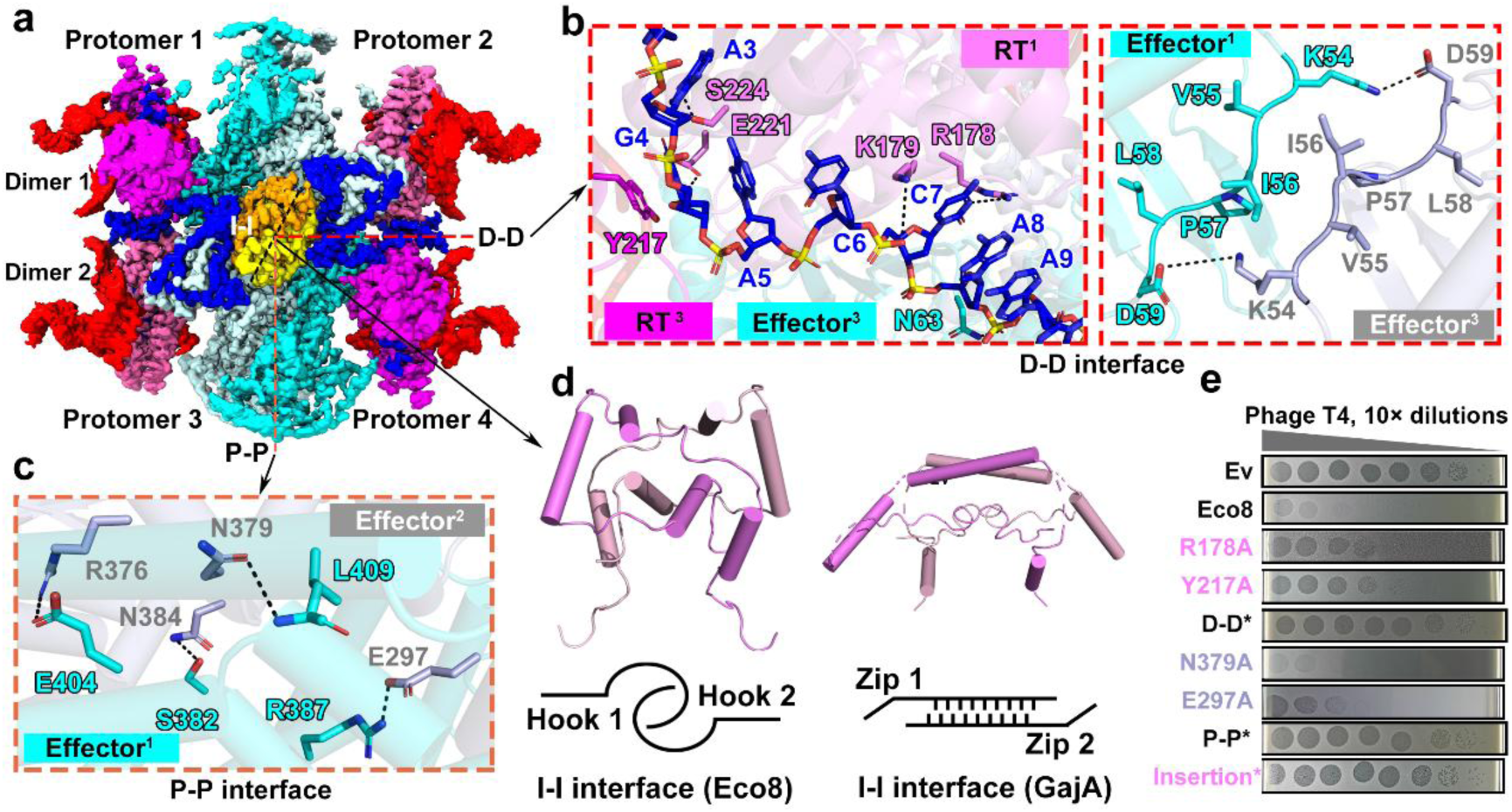
Retron-Eco8 tetramer formation mediated by both inter- and intra-interactions. **a** Surface representation of the Retron-Eco8^ATP^ structure. The D-D, P-P and I-I interfaces are labeled with dashed lines. **b, c, d** Detailed views for D-D interface (b), P-P interface (c) and I-I interface (d) (GajA I-I interface: PDB, 8SM3). **e,** Serial dilution plaque assays for T4 phage on *E. coli* MG1655 transformed with plasmids encoding WT or indicated interface mutants of RT and effector proteins. (D- D*: R178A/K179A/Y217A/E221A; P-P*: R376A/N384A/R387A; Insertion*:170N- 251T→GSGSGSGSGSG).

The nucleic acid topology closely resembles that of other characterized retrons, comprising an msrRNA (rG18–rG81) and a reverse-transcribed msdDNA (dC1– dT75)^13,14^. These nucleotides form a partially pairing central duplex, flanked by two inverted repeats IRa2 and IRa1. These inverted repeats are bordered by two conserved guanosine residues—the branching G and the opposing G—that form a short dsRNA element (Fig. 2c). In our structure, the 3′ end of the msrRNA and both ends of the msdDNA are clearly resolved, allowing us to estimate the lengths of msrRNA and msdDNA as 64 and 75 nucleotides, respectively (Fig. 2c), consistent with prior biochemical characterizations^2^. Together, these data reveal that Retron-Eco8 assembles into a tetrameric complex, in which each protomer contains an RT, the associated msrRNA–msdDNA hybrid, and an effector, forming the structural basis for its anti-phage function.

### Mechanistic insights into msdDNA synthesis and termination by the RT subunit

The msdDNA is reverse transcribed at the active site of the RT subunit, using the *msd* region of the ncRNA as a template. Structural analysis identified several conserved motifs critical for single-stranded DNA synthesis, including the YADD (Y198–D201) and ING (I247–G249) motifs in the palm domain, and the PMG (P163–G165) motif in the fingers domain (Fig. 2d and Extended Data Fig. 3e). A coordinated Mg²⁺ ion, positioned by the aspartate residue D107 and the YADD motif, likely facilitates catalysis. In addition, the PMG motif and a conserved phenylalanine residue (F112) form a pocket that constrains the terminal deoxynucleotide (Fig. 2d). Alanine substitution of residues in the YADD, ING, and PMG motifs, as well as D107 and F112, abolished anti-phage defense activity (Fig. 2e), highlighting the essential role of reverse transcription in retron-mediated immunity.

A key mechanistic question is how reverse transcription is precisely terminated at nucleotide A73 of the msrRNA. Structural analysis revealed that the terminal adenine A73, which base-pairs with the last deoxynucleotide T75, is stacked against residue F51 in the β-hairpin. This stacking interaction induces a ∼90° bend in the preceding nucleotide (U72), sterically blocking its entry into the catalytic site and thereby terminating DNA synthesis (Fig. 2f). Collectively, these findings elucidate the molecular basis of msdDNA synthesis and precise termination by Retron-Eco8.

### Tetramerization is essential for the anti-phage activity of Retron-Eco8

The Retron-Eco8 complex adopts a tetrameric architecture, consisting of two tightly associated dimers (dimer 1: protomers 1 and 2; dimer 2: protomers 3 and 4). Two dimers are firmly connected together through dimer-to-dimer (D-D) interface (Fig. 3a, b). This D-D interface is stabilized by both protein-protein interactions (Fig. 3b, right inset) and protein-msdDNA interactions (Fig. 3b, left inset). The msdDNA acts as a bridging scaffold, forming specific interactions with adjacent RT protomers (Fig. 3b, left inset).

Within each dimer, protomer 1 and 2 (protomer 3 and 4) associate exclusively via their effector proteins, forming a protomer-to-protomer (P-P) interface (Fig. 3a, c). In details, this interface is stabilized by a 2-fold symmetric network of hydrogen bonds (S382/L409 to N384/N379) and salt bridges (R387/E404 and E297/R376) (Fig. 3c). Further stabilization of the tetramer arises from insertion-to-insertion (I-I) interfaces between protomers 1–4 and 2–3 (Fig. 3a). Unlike the GajA complex, where insertions form a central α-helical "zipper," Retron-Eco8 insertions adopt a hook-like structure that grasps diagonal protomers, collectively occupying the central pore (Fig. 3d). Alanine substitutions of key residues in the D-D (R178A/K179A/Y217A/E221A) and P-P (R376A/N384A/R387A) interfaces, or replacement of the insertion with a GS linker, completely eliminated the phage defense activity *in vivo* (Fig. 3e), highlighting the importance of these interactions in maintaining the overall structure of Retron-Eco8.

### Phage SSB directly binds msdDNA and activates the Retron-Eco8 system

Previous studies showed that single-stranded DNA-binding proteins (SSBs) from phages SECphi4 bind the msdDNA of Retron-Eco8, thereby triggering system activation and cellular toxicity *in vivo*^2,3^. Consistently, co-expression of Retron-Eco8 with the SSB from T7 phage—which is sensitive to Retron-Eco8 (Fig. 1c)—activated the system and induced cell death (Fig. 4a). To determine whether phage T7 SSB can directly bind to msdDNA *in vitro*, we purified phage T7 SSB and incubated it with the Retron-Eco8 complex. Pull-down assays confirmed their direct interaction (Fig. 4b). Deletion of two predicted SL structures within msdDNA abolished this interaction, without affecting overall Retron-Eco8 assembly (Fig. 4b and Extended Data Fig. 4a), suggesting that SSB specifically binds these SLs. Further cryo-EM analysis of the Retron-Eco8–SSB complex revealed two rows of additional globular densities decorating the outer surface of the effector (Fig. 4c). Although these densities are conformationally flexible and visible only at low contour levels, precluding confident fitting with the AF3-predicted SSB model, their spatial coincidence with the msdDNA position—together with our biochemical evidence—strongly supports direct SSB engagement with the msdDNA of Retron-Eco8.

**Fig. 4.**
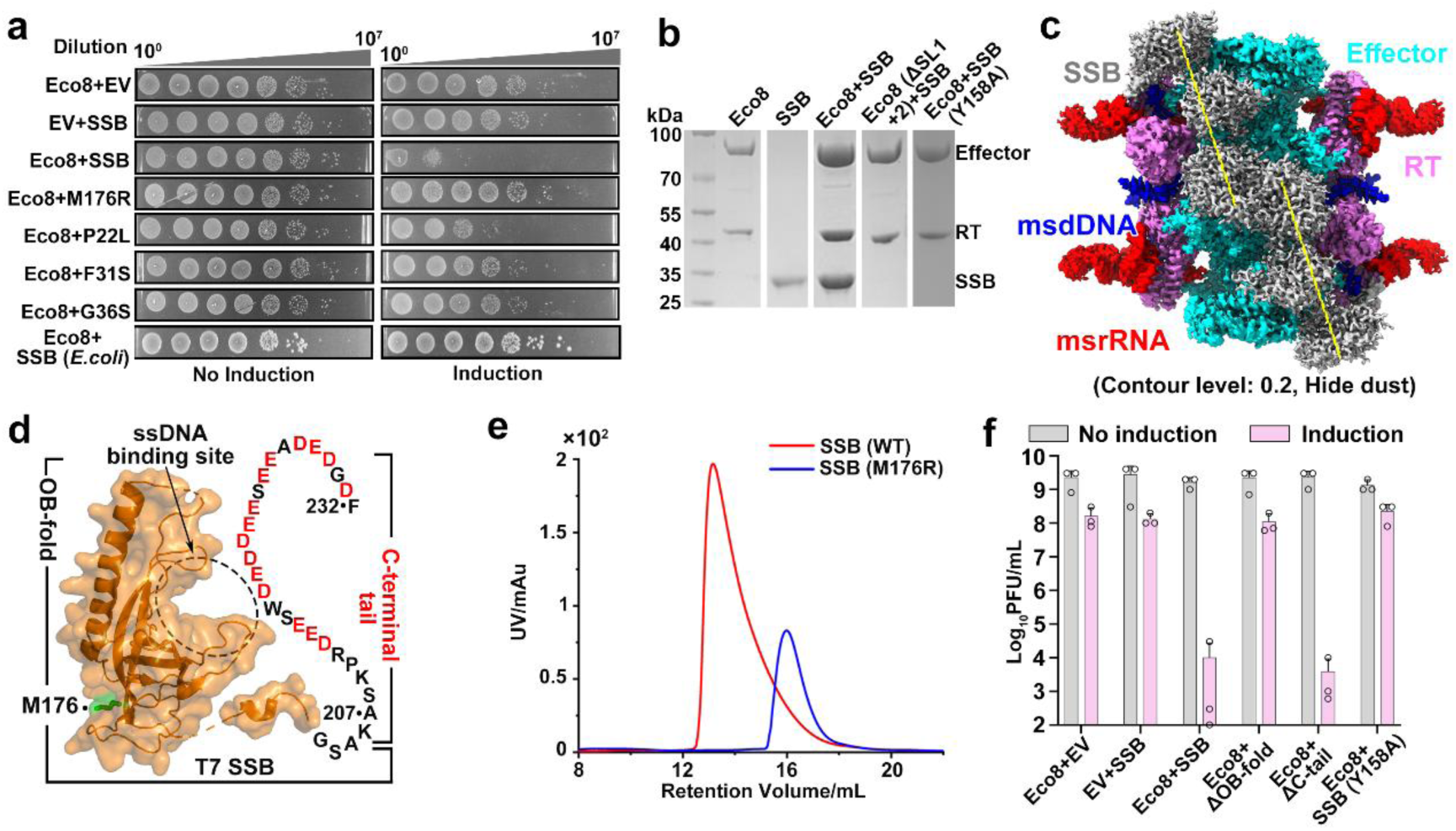
Phage T7 SSB binds msdDNA to activate the Retron-Eco8 immune system. **a, f** Survival status of *E. coli* cells co-producing Retron-Eco8 and *E.coli* or T7 SSB (WT or mutants). Data are representative of three biological replicates. (n = 3; mean ± SD). **b** *In vitro* pull-down assay using Strep II-tagged Retron-Eco8 (WT or SL truncation) to capture SSB (WT or Y158A mutant). **c** Surface representation of the cryo-EM structure of the Retron-Eco8-SSB complex. For better presentation, the surrounding densities present in the Retron-Eco8 complex have been omitted. **d** Surface view of the phage T7 SSB (PDB, 5ODL), the disordered C-terminal tail is shown, with acidic residues labeled in red. The residue M176 is colored in green and the ssDNA binding site is highlighted by dashed-line box. **e** Size exclusion chromatography profiles of T7 SSB (WT) and (M176R) mutant.

### Oligomerization and DNA binding ability of phage-derived SSB are essential for Retron-Eco8 activation

The direct binding of phage SSB to the msdDNA of Retron-Eco8 activates the system, raising the question of whether the host-derived SSB can do the same. Although *E. coli* SSB forms a homotetramer capable of binding ssDNA^15^, co-expression with retron- Eco8 demonstrated its inability to trigger system activation (Fig. 4a), indicating that activation requires a specific phage-derived SSB. Previous studies identified a T7 SSB mutant (M176R) that escapes Retron-Eco8-mediated toxicity^3^. However, residue M176 is positioned on the outer surface of T7 SSB, distant from the ssDNA binding site (Fig. 4d)^16^. Since phage SSB proteins typically function as oligomers to facilitate high-affinity binding to ssDNA during phage replication^17^, we hypothesized that M176R mutant might affect SSB oligomerization, thereby impairing Retron-Eco8 activation. To test this hypothesis, we performed size-exclusion chromatography (SEC) with purified wild-type T7 SSB and M176R mutant. The M176R mutant exhibited a delayed elution profile relative to the wild-type protein (Fig. 4e), consistent with a disruption of its oligomeric state. To further assess the role of SSB oligomerization in Retron-Eco8 activation, we tested three additional mutations (P22L, F31S, and G36S) in T7 SSB, which is known to impair dimerization (Extended Data Fig. 4b)^18^. All of these mutants failed to activate Retron-Eco8 to induce cellular toxicity (Fig. 4a). These results indicate that the oligomeric state of phage SSB is critical for system activation.

Given that phage SSB physically interacts with Retron-Eco8, we next sought to identify the specific region responsible for this interaction. T7 SSB comprises an N-terminal oligosaccharide/oligonucleotide-binding fold (OB-fold, residues 1–206) and a short C-terminal acidic tail (C-tail, residues 207–232) (Fig. 4d). While the OB-fold mediates ssDNA recognition, the C-tail has been implicated in protein–protein interactions with key T7-encoded proteins, including DNA polymerase, helicase-primase and other factors essential for T7 DNA metabolism^17^. Deletion of the C-tail did not affect Retron-Eco8 activation, whereas removal of the OB-fold completely abolished activation (Fig. 4f), underscoring its essential role. To identify residues within the OB-fold required for Retron-Eco8 activation, we aligned the T7 SSB structure with its homolog from phage Enc34 in complex with ssDNA¹² (Extended Data Fig. 4c). This comparison highlighted a conserved aromatic residue, Y158, predicted to contact nucleic acids. Substitution residue Y158 with alanine abolished Retron-Eco8 activation (Fig. 4f) and eliminated SSB interaction with the Retron-Eco8 complex in pulldown assays (Fig. 4b). These findings demonstrate that phage SSB directly binds to Retron- Eco8 msdDNA to trigger system activation, with its oligomeric state and nucleic acids-binding ability being indispensable for this process, highlighting the requirement for phage-derived SSB in Retron-Eco8 activation.

### Phage SSB-activated Retron-Eco8 degrades both phage and host DNA

The efficient activation of Retron-Eco8 by phage T7 SSB to induce cell death prompted us to investigate the underlying mechanism of toxicity. The Retron-Eco8 effector belongs to the OLD family of nucleases, which are metal-dependent endonucleases known to mediate potent DNA cleavage activity *in vitro*^19,20^. This phylogenetic and biochemical features suggest that the Retron-Eco8 effector may exert its toxic activity through DNA degradation upon system activation. To test this hypothesis, we used fluorescence microscopy to visualize nucleic acid integrity in *E. coli* MG1655 cells co-expressing wild-type or mutant Retron-Eco8 alleles together with phage T7 SSB following 4 hours of induction. Cells were stained with DAPI to visualize DNA and FM4-64 to label membranes. A substantial proportion of cells expressing the wild-type Retron-Eco8 system showed markedly decreased DAPI fluorescence, indicating DNA degradation (Fig. 5a). In contrast, cells expressing catalytically inactive variants of the reverse transcriptase (YADD to YAAA) or the Toprim effector (D450A/D452A) retained normal DNA staining (Fig. 5a). These observations suggest that Retron-Eco8 triggers cell death through the nonspecific nuclease activity of its Toprim effector, likely by degrading host DNA.

**Fig. 5.**
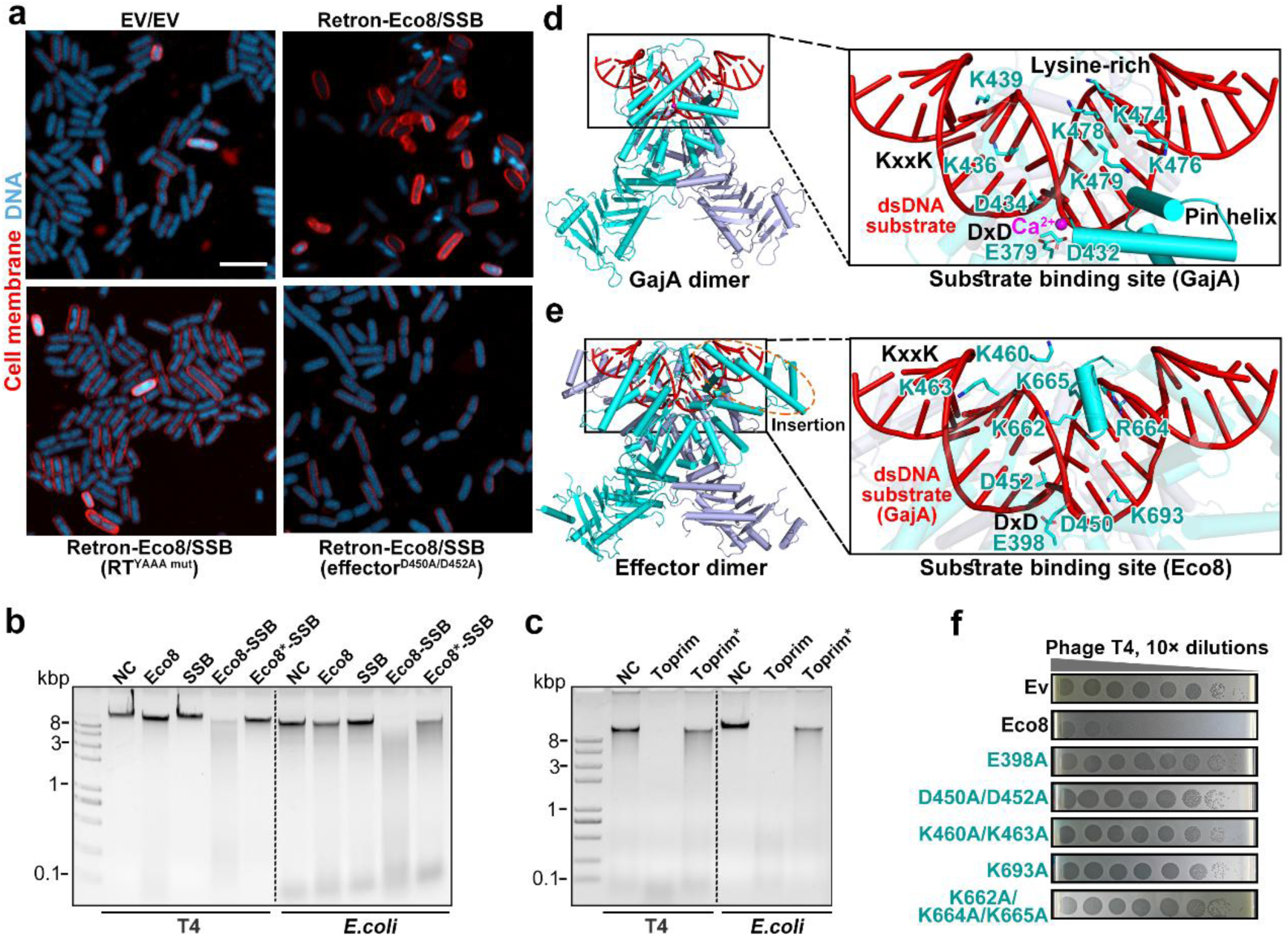
Phage SSB binding activates Retron-Eco8 system to degrade intracellular DNA. **a** Representative fluorescence microscopy images of *E.coli* MG1655 expressing the Retron-Eco8 (WT, RT^YAAA^ mutant or effector^D450A/D452A^ mutant) and T7 SSB protein. Red = outer membrane (FM4-64 stained), cyan = DNA (DAPI stained). Scale bar = 5 μm. Imaging was performed in biological triplicate. **b, c** *In vitro* nuclease activity assays of Retron-Eco8-SSB complex (b) or Toprim domain alone (c) against *E. coli* and phage T4 genomic DNA. Eco8*-SSB and Toprim* represent D450A/D452A mutants. **d, e** Overall (left) and detailed (right) views of the GajA dimer (8JQ9) (d) and effector protein of Retron-Eco8 complex superimposed with dsDNA substrate (e). Key residues involved in substrate recognition are labeled. **f** Serial dilution plaque assays for T4 phage on *E. coli* MG1655 strain transformed with plasmids encoding WT or indicated effector protein mutants.

To further examine whether phage SSB activates Retron-Eco8 to degrade phage DNA and the *E. coli* genome *in vitro*, we incubated the Retron-Eco8–SSB complex with either phage T4 or *E. coli* genomic DNA. The results showed that Retron-Eco8–SSB directly degraded both substrates, whereas mutation of the key residues (D450A/D452A) within the Toprim domain of the effector protein substantially impaired its DNase activity (Fig. 5b). Notably, Retron-Eco8 lacking phage SSB showed no DNase activity, further confirming that it adopts an autoinhibited conformation (Fig. 5b). Consistently, a truncated Toprim domain retained DNase activity against both phage and host DNA independent of phage SSB (Fig. 5c), supporting that full-length Retron-Eco8 is autoinhibited until its nuclease activity was relieved by phage SSB binding. Together, these findings indicate that phage SSB directly interacts with Retron- Eco8 to activate its Toprim’s nuclease activity, leading to degradation of both phage and host DNA, consistent with our *in vivo* observations.

### Mechanistic insights into Retron-Eco8 activation and effector function

To define the DNase active site of Retron-Eco8, we aligned the dsDNA-bound GajA structure with the effector protein of Retron-Eco8, and found that dsDNA substrate from GajA fits well into the catalytic channel of the effector (Fig. 5d, e). In GajA, the dsDNA substrate is clamped between two Toprim domains through three primary interfaces—the KxxK, lysine-rich, and Pin-helix motifs (Fig. 5d)^12^. Retron-Eco8 retains the conserved DxD catalytic residues (D450, D452) and the KxxK motif (K460- K463) (Fig. 5e), and mutation of either features abolished its anti-phage activity (Fig. 5f). Notably, the Retron-Eco8 Toprim domain is larger than that of GajA and contains an additional all α-helical insertion (residues 568–638 and 651–696) that contributes to form the substrate-binding channel (Fig. 5d, e). This insertion is enriched in positively charged residues (K693, K662, R664, K665) (Fig. 5e), and alanine substitutions at these positions substantially reduced its phage defense activity (Fig. 5f), highlighting the importance of substrate recognition for nuclease function. Together, these findings define the DNA-binding sites within the Toprim domain of Retron-Eco8.

A key mechanistic question is how phage SSB binding to the SLs of msdDNA and activates its nuclease activity. OLD-family nucleases typically require conformational changes in their DNA-binding channels to widen and accommodate dsDNA substrates^12^. However, comparison of the Retron-Eco8 and Retron-Eco8–SSB complexes revealed no significant structural changes in the Toprim domain, likely owing to the low occupancy of SSB or the absence of bound DNA in the cryo-EM reconstruction. To reveal potential conformational transitions upon DNA engagement, we predicted the structure of the effector dimer bound to dsDNA using AlphaFold3^21^. The model revealed a marked expansion of the substrate-binding channel, with the distance between the innermost α-helices increasing from 14.8 Å to 17.8 Å, as measured between the Cα atoms of K665 in adjacent protomers (Fig. 6a, top inset). Another prominent change involves the FDYY loop (residues 481–484), which shifts downward by ∼10 Å upon activation (Fig. 6a, bottom inset). In the autoinhibited Retron-Eco8 complex, this loop interacts directly with the msdDNA–msrRNA duplex, where Y483 and Y484 form π–π stacking interactions with nucleotides G70 and A69, respectively (Fig. 6b). Mutation of the FDYY motif to AAAA severely impaired phage defense activity (Fig. 6c), underscoring the essential role of this loop in Retron-Eco8 activation.

**Fig. 6.**
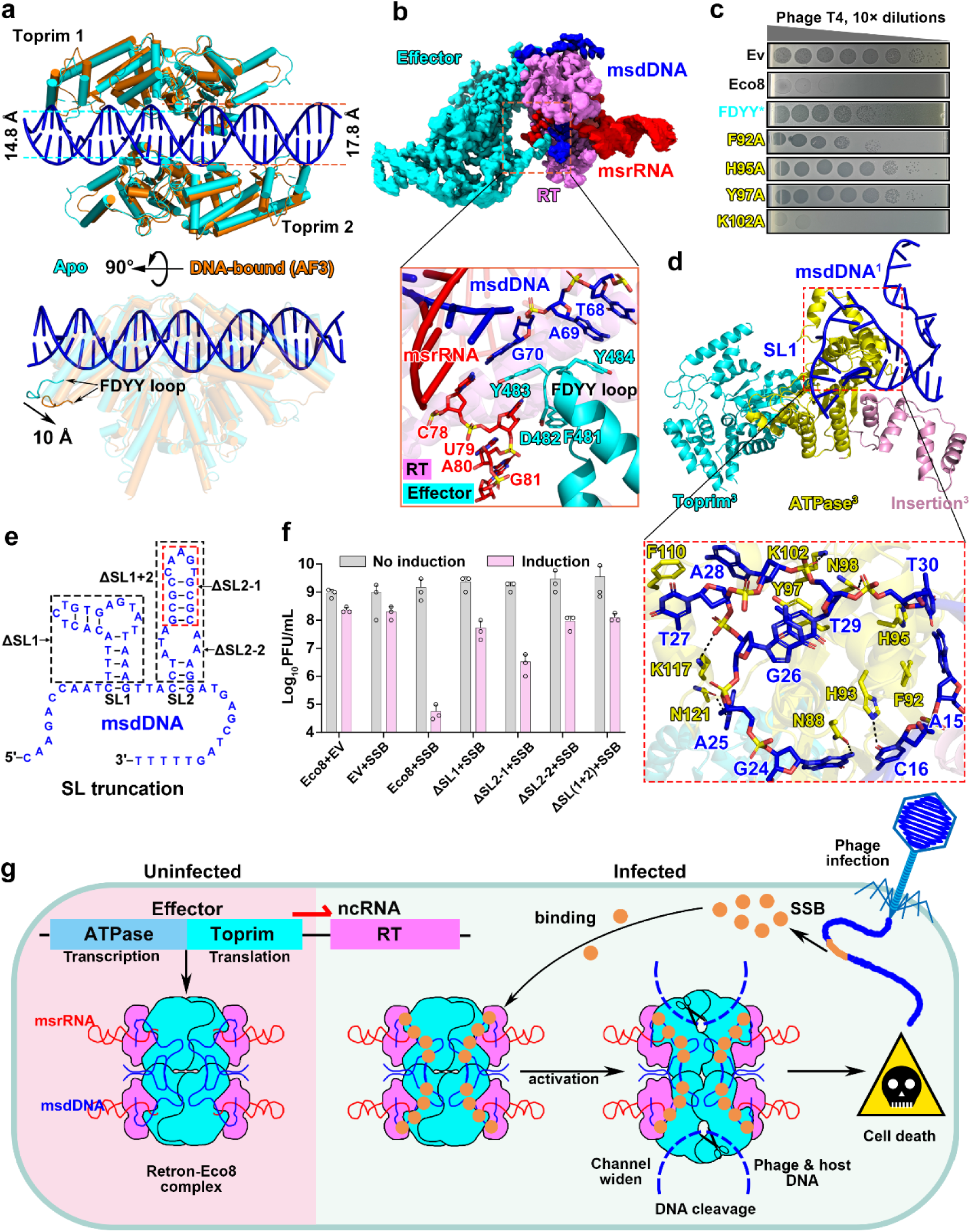
Molecular basis of phage-SSB-mediated Retron-Eco8 activation **a** Structural comparison of the Toprim dimers of Retron-Eco8 apo (cyan) and AF3- predicted DNA-bound (30 bp) states (orange). **b** Overall surface (above) and detailed ribbon (below) representations of one Retron- Eco8 protomer. **c** Serial dilution plaque assays for T4 phage on *E. coli* MG1655 strain transformed with plasmids encoding WT or indicated mutants. (FDYY*:481FDYY485/481AAAA485). **d** The effector protein from protomer 3 is encircled by SL from protomer 1. Key residues responsible for interactions are labeled and hydrogen bond interactions are labeled with dashed line. **e** Schematic diagram shows the different truncations of the SL region of the msdDNA. **f** Survival status of *E. coli* MG1655 cells co-producing Retron-Eco8 (WT or SL truncations) and T7 SSB. Data are representative of three biological replicates. (n = 3; mean ± SD). **g** Mechanistic model for activation and effector function of bacterial Retron-Eco8 immunity.

Next we sought to determine which region of T7 SSB binds to msdDNA for Retron- Eco8 activation. As truncation of the msdDNA SL structure abolished its direct interaction with T7 SSB, this indicates that SL mediates SSB recognition. In our structure, a 5-nt bubble (A25–T29) within SL1 from protomer 1 forms extensive π–π stacking and hydrogen-bonding interactions with the ATPase domain of protomer 3, partitioning SL1 into two orthogonal segments (Fig. 6d). Deletion of SL1 (ΔSL1) or point mutations disrupting the SL1–ATPase interface (F92A, H95A, and Y97A) impaired both SSB-induced activation and phage resistance (Fig. 6c, e, f), highlighting its crucial role in Retron-Eco8 activation. The rigid configuration of SL1 likely precludes SSB engagement, whereas the absence of electron density corresponding to SL2 suggests a flexible conformation, suggesting that SL2 serves as the potential SSB-binding site. Partial deletion of SL2 (ΔSL2-1) markedly reduced SSB-dependent activation, while complete removal of SL2 (ΔSL2-2) abolished the activation (Fig. 6e, f), demonstrating that SL2 is essential for SSB-mediated activation. Together, these findings indicate that SL structures are crucial for Retron-Eco8 activation, with T7 SSB binding to SLs likely disrupting msdDNA–effector interactions, releasing autoinhibition of the Toprim domain, and thereby enabling nuclease activation.

## Discussion

Retrons constitute a diverse class of bacterial defense systems distinguished by their effector modules, which range from OLD-family endonucleases and TIR domains to proteases, transmembrane proteins, and cold-shock factors^2^. Retron-Eco8, composed of a reverse transcriptase (RT), a noncoding RNA, and an OLD-family nuclease effector, provides antiphage immunity by triggering abortive infection upon sensing a phage-encoded single-stranded DNA-binding (SSB) protein^3^. Our structural and functional analyses elucidate the molecular basis underlying system activation and subsequent effector function. Retron-Eco8 assembles into a heteromeric complex with a stoichiometry of 4:4:4 (RT: effector: msDNA). Within this architecture, the effector subunits form a central scaffold encircled by RT subunits, which are further wrapped by msdDNA. This configuration locks the effector in an autoinhibited state. Upon phage infection, phage-encoded SSB proteins bind the stem-loop structures of msdDNA, relieving msdDNA-mediated autoinhibition, likely promoting widening of the double-stranded DNA–binding channel to activate the effector nuclease domain. The activated nuclease then catalyzes non-specific degradation of intracellular DNA, including both phage and host genomes, leading to host cell death that prevents phage propagation (Fig. 6g).

These findings, together with previous studies^2–4,13,14,22–26^, indicate that msdDNA acts as both an autoinhibitor and a molecular sensor. In Retron-Eco8, msdDNA encircles the effector core, enforcing an inactive conformation. The self-inhibitory function of msdDNA appears to be a mechanism shared by several retron systems. For instance, in the Retron-Eco1 system, msDNA sequesters the effector dimer into a filamentous assembly, thereby neutralizing its toxicity^13^. Retron-Ec83–Septu employ an ATPase-associated HNH nuclease domain to cleave single-stranded DNA^23^, while Retron-Ec78 utilizes the same effector as Retron-Ec83-Septu but targets tRNA for cleavage^22,24–26^. In both systems, effector activity is suppressed through assembly into an autoinhibited complex with RT and msDNA. As a sensor, Retron-Eco8 activation is initiated by direct recognition of msdDNA by phage-derived SSB, paralleling other retron systems that respond to DNA-interacting enzymes. Retron-Sen2, for instance, is activated by the host DNA adenine methyltransferase (Dam)^27^, Retron- Eco1 senses cytosine methylation of its msDNA by a phage-encoded DNA methyltransferase (Dcm)^14^, and Retron-Eco11 is activated by helicases from phages T4 and T5^28^. Together, these examples highlight a surveillance strategy in which retrons monitor perturbations to msdDNA as a cue for phage infection. Beyond their defensive role, the retron msd sequence exhibits considerable variability and flexibility, allowing incorporation of diverse inserts that direct intracellular synthesis of corresponding single-stranded DNA^6,7^. This property underlies the repurposing of retron-derived ssDNA as templates for recombineering and genome engineering across both prokaryotic and eukaryotic systems^29^. The flexibility of Retron-Eco8 msdDNA suggests that it may likewise serve as a platform for genome-editing applications.

The Retron-Eco8 effector belongs to the OLD-family nucleases, which contain both an ATPase and a Toprim nuclease domain. While the Toprim domain mediates DNA cleavage, the role of the ATPase domain remains enigmatic. In our cryo-EM reconstruction of Retron-Eco8 in the presence of 5 mM ATP, no ATP density was observed despite the conservation of ATP-binding motifs. Consistently, the purified Retron-Eco8 complex exhibited no measurable ATPase activity in vitro, regardless of the presence of SSB or dsDNA (Extended Data Fig. 5a). Nevertheless, alanine substitutions in key residues within the ATP-binding pocket abolished antiphage activity in vivo (Extended Data Fig. 5b, c), indicating that ATP-binding motifs are essential for system function. Given that high ATP concentrations suppress the nuclease activity of the Gabija OLD-family defense system^12^, we examined whether ATP exerts a similar effect on Retron-Eco8 activation. Indeed, nuclease activity was markedly inhibited at physiologically relevant ATP levels^30^ (≥ 3 mM) (Extended Data Fig. 5d), suggesting that Retron-Eco8 is likewise subject to ATP-mediated regulation. These findings point to a critical regulatory role for ATP in controlling Retron-Eco8 activity, although the underlying molecular mechanism remains to be elucidated.

Taken together, our findings, in parallel with recent two studies^31,32^, delineate the molecular mechanism of Retron-Eco8 complex assembly, autoinhibition, phage SSB– mediated activation, and effector function. These insights deepen our understanding of retron-mediated antiphage immunity and provide a structural foundation for future retron-based biotechnological applications.

## Methods

### Bacterial strains

*Escherichia coli* DH5α, BL21-Star (DE3), BL21-AI and *MG1655* cells were used for plasmid construction, protein expression, *in vivo* plasmid interference assay and phage interference assay, respectively.

### Protein expression and purification

The full-length Retron-Eco8 sequences of *Escherichia coli* 200449 was synthesized by SYNBIOB GENE and cloned into the pRSFDuet-1vector (Novagen). The construct was designed to fuse an N-terminal 6×His tag to the effector protein and a C-terminal Strep-II tag to the reverse transcriptase (RT) protein. All constructs were verified by Sanger-sequencing and expressed in *E. coli* BL21-Star (DE3) cells. For protein expression, a single colony was inoculated into LB medium supplemented with 50 μg/mL kanamycin and grown overnight at 37 °C. The culture was then diluted into fresh LB medium and cultured at 37°C until the OD600 reached 1.0. Protein expression was induced by 0.5 mM isopropyl-β-D-1-thiogalactopyranoside (IPTG), followed by incubation at 16 °C for 16 h.

Cells were harvested by centrifugation and resuspended in a lysis buffer (25 mM Tris-HCl, 150 mM NaCl, 5 mM imidazole, pH 8.0). The cell suspension was sonicated on ice for 40 min and centrifugation at 16,000 × g for 30 min at 4 °C. The supernatant was loaded onto a 5 mL His Trap Fast Flow column (Cytiva Life Sciences) and allowed to bind by circulating the lysate through the column for 1 hour at 4 °C. The column was washed with 10 column volumes (CV) of lysis buffer, and bound proteins were eluted with 20 mL of elution buffer (25 mM Tris-HCl, 150 mM NaCl, 300 mM imidazole, pH 8.0). The eluate was then applied to a Strep-tactin Sepharose column pre-equilibrated with Buffer A (25 mM Tris-HCl, 150 mM NaCl, pH 8.0). After washing with 10-15 CV of buffer A, Retron-Eco8 was eluted with buffer A containing 2.5 mM D-desthiobiotin (Smart-lifesciences). The eluted sample was diluted with Q binding buffer (25 mM Tris-HCl, 50 mM NaCl) and loaded onto a 5 mL Q column (Cytiva Life Sciences) pre-equilibrated with the same buffer. Proteins were eluted by a linear gradient of 50 mM to 1 M NaCl over 20 CV. Target fractions were concentrated using a 50 kDa molecular weight cut-off concentrators (Merck Millipore, UFC903024) and further purified by size-exclusion chromatography on a Superdex 200 Increase 10/300 GL column (Cytiva Life Sciences) pre-equilibrated with running buffer (20 mM Tris-HCl, 150 mM NaCl, pH 7.5). The pooled Retron-Eco8 fractions were pooled, concentrated, flash-frozen in liquid nitrogen, and stored at -80°C for subsequent experiments.

For the Retron-Eco8^ATP^ complex, which was constructed with a C-terminal Strep- II tag, the purification procedure was performed as described above.

All mutants were generated through site-directed mutagenesis and purified by the same method as described above.

### Cryo-EM sample preparation and data acquisition

3.0 μL of a ∼16 mg/mL solution of purified Retron-Eco8 or Retron-Eco8-SSB complexes were applied onto glow-discharged UltrAuFoil 300 mesh R1.2/1.3 grids (Quantifoil). The grids were blotted for 2 s at 100% humidity and flash frozen in liquid ethane using a FEI Vitrobot Mark IV (FEI). Images were collected on a Titan Krios electron microscope (FEI) operated at an acceleration voltage of 300 kV, equipped with a Gatan K3 Summit detector with a physical pixel size of 1.1 Å. The defocus during image collection ranged from -1.0 µm to -2.5 µm. Images were dose-fractionated into 32 frames with a total accumulated dose of 50 electrons per Å^2^.

To obtain the Retron-Eco8^ATP^ complex for structural determination, the purified Eco8 protein was equilibrated on ice for 30 minutes in a buffer containing 50 mM Tris-HCl (pH 8.0), 5 mM MgCl₂, and 10 mM ATP. The sample was then concentrated and prepared using the same methods as previously described.

### Cryo-EM data processing

Image processing was conducted by RELION 3.1^33^ and cryoSPARC v3.1^34^. Motion correction was performed with MotionCor2^35^. Contrast transfer function (CTF) parameters were estimated by Ctffind4^36^. Auto-picked particles using the Laplacian-of-Gaussian method were extracted and underwent two rounds of 2D classification to remove junk particles and several rounds of heterogeneous refinement in cryoSPARC v3.1, utilizing an initial model generated within cryoSPARC v3.1 as a reference.

Particles corresponding to the best class with the highest-resolution features were selected and subjected to non-uniform refinement in cryoSPARC v3.1. These particles then underwent CTF refinement followed by another round of non-uniform refinement, generating the final reconstitutions. All resolutions were estimated by applying a soft mask around the protein density and the Fourier shell correlation (FSC) = 0.143 criterion in cryoSPARC v3.1. Local resolution estimates were calculated from two half data maps in cryoSPARC v3.1. Further details related to data processing and refinement are summarized in Supplementary Table 1.

### Model building and refinement

All atomic models were built with the help of AF predicted structure, and then manually refined by COOT v0.9.5^37^ interactively. All models were then refined against combined maps using phenix.real_space_refine v1.19.2^38^ by applying geometric and secondary structure restraints. All structural figures were prepared in Pymol v2.4.2 (http://www.pymol.org) and Chimera X^39^.

### Cell viability analysis assay

*E. coli* BL21-AI cells were transformed via heat-shock with pETDuet1 plasmids encoding either Retron-Eco8 or its variants, together with pBAD (ori: pBR322) plasmids encoding either T7-SSB or its variants. Transformed cells were grown overnight at 37 °C in LB medium supplemented with 2% glucose and appropriate antibiotics (50 μg/mL kanamycin, 50 μg/mL ampicillin). The culture was then diluted 1:100 into fresh LB medium containing 2% glucose and antibiotics, and incubated at 37 °C until OD600 reached 0.6. Cells were harvested, washed, and resuspended in glucose-free LB medium. Serial dilutions of the resuspended cells were prepared and spotted onto LB agar plates containing antibiotics along with either inducers (1% L-arabinose and 1 mM IPTG) or 2% glucose as a control.

For cell viability assays involving Retron-Eco8 SL truncations, *E. coli* MG1655 cells were transformed by heat-shock with a pACYC184 plasmid encoding either wild-type Retron-Eco8 or its SL truncations, along with a pBAD plasmid expressing T7-SSB.

### Phage plaque assays

For Phage plaque assays, *E. coli* MG1655 transformed with pACYC184 plasmids (containing WT or mutants) was grown overnight at 37 °C with shaking at 220 rpm After overnight incubation, 300 μL of the bacterial culture was mixed with 30 mL molten MMB agar (LB, 5 mM MgCl2, 5 mM CaCl2, 0.1 mM MnCl2, 50 μg/mL tetracycline, 0.5% agar). The mixture was poured onto square plates (biosharp) and dried for 60 min at room temperature. Tenfold serial dilutions in phosphate buffered saline (PBS) were prepared for each of tested phages. Once the top agar solidified, 10 μL drops of the diluted phage culture were spotted on the bacterial layer. Plaque-forming units (PFUs) were determined by counting the derived plaques after overnight incubation. Phage defense activity was assessed by calculating the fold reduction in efficiency of plating (EOP), which was determined by dividing the plaque-forming units per ml (PFU/mL) obtained on a lawn of empty vector (EV) control cells by the PFU/mL obtained on a lawn of defense system-expressing cells.

### Nuclease activity assays

Nuclease activity was assessed in 10 μL reactions containing 150 ng of *E. coli* genomic DNA or phage T4 genomic DNA, and 1 μM of the respective protein in reaction buffer (20 mM Tris-HCl, pH 8.0, 5 mM MnCl₂). Reactions were incubated at 37°C for 90 min, followed by treatment with 1 μL Proteinase K to degrade proteins. The reactions were terminated by adding 1 μL of 10 × loading dye, and the products were resolved on 1% native agarose gel.

For assays involving varying ATP concentrations, ATP solutions were pH-adjusted by mixing 50 μL of 0.1 M ATP with 50 μL of 1 M Tris-HCl (pH 8.0), followed by dilution to the desired concentrations.

### Fluorescence microscopy

Overnight cultures of *E. coli* MG1655 carrying Retron-Eco8 (pACYC184) and T7- SSB (pBAD) or empty vectors were diluted 1:100 and grown at 37°C to an OD600 of 0.5–0.6 in the presence of kanamycin (50 μg/mL), tetracycline (50 μg/mL), and 2% glucose. The cells were then resuspended in fresh medium containing antibiotics and 1% L-arabinose and incubated for an additional 4 h at 37 °C. Prior to imaging, samples were stained with DAPI (4 μg/mL, 37 °C, 30 min) and FM4-64 (2 μg/mL, 4 °C, 1 min), and visualized by fluorescence microscopy.

### ATPase assays

ATP hydrolysis activity was measured using a commercial phosphate detection kit by quantifying free inorganic phosphate release. Reactions (50 μL) contained 2 mM ATP, 100 ng of 5,000-bp dsDNA and 2 μM proteins in reaction buffer (20 mM Tris-HCl pH 8.0, 100 mM NaCl, 5 mM MgCl2). After incubation at 37 °C for 2 h, absorbance was measured at 630 nm using an Agilent BioTek Synergy H1 microplate reader.

### Cell survival assay

Overnight cultures of *E. coli* MG1655 expressing WT Retron-Eco8 (pACYC184) or an empty vector control were diluted 1:100 and grown at 37°C until reaching an OD600 of 0.5–0.6 in LB medium supplemented with 5 mM MgCl2, 5 mM CaCl2, 0.1 mM MnCl2, 50 μg/mL tetracycline. Cells were infected with T4 phage at multiplicities of infection (MOI) of 0, 0.5, and 5. Samples were collected before infection and at 2, 15, 30, and 60 min post-infection. Serial 10-fold dilutions of each sample were prepared, and 6 μL aliquots were spotted onto LB agar plates containing tetracycline (50 μg/mL). Colonies were counted after overnight incubation at 37 °C.

## Data Availability

The cryo-EM density maps have been deposited in the Electron Microscopy Data Bank (EMD-62954 for Retron-Eco8 complex [https://www.ebi.ac.uk/emdb/EMD-62954], EMD-66110 for Retron-Eco8^ATP^ complex [https://www.ebi.ac.uk/emdb/EMD-66110], and EMD-63117 for Retron-Eco8-SSB complex [https://www.ebi.ac.uk/emdb/EMD-63117]). The atomic coordinates have been deposited in the Protein Data Bank (PDB) with accession number 9LBQ (Retron-Eco8 complex) [https://www.rcsb.org/structure/9LBQ] and 9WN8 (Retron-Eco8^ATP^ complex) [https://www.rcsb.org/structure/9WN8]. This paper does not report original code. Source data are provided with this paper.

## Acknowledgements

We thank the staff at Southern University of Science and Technology (SUSTech) Cryo EM Center for assistance in data collection on the SUSTech Titan KRIOS cryo-electron microscope. This work was funded by the National Key R&D Program of China (2024YFE0198900); National Natural Science Foundation of China (grant no. 32400024 to J.T.Z.); Shenzhen Key Laboratory of Prevention and Treatment of Severe Infections (grant No. ZDSYS20200811142804014 to H.Y.L.); Shenzhen Natural Science Foundation (grant No. JCYJ20240813094609013 to J.T.Z.). N.J. is an investigator of SUSTech Institute for Biological Electron Microscopy.

## Author contributions

N.J., X.L. and J.-T.Z. conceptualized the strategy. C.-G.J., Z.L. and Y.-L.L. .-performed the in vitro and in vivo experiments. Cryo-EM sample preparation, data collection, processing and analysis, structure refinement are performed by J.-T.Z. and X.-Y.W. N.J. and J.-T.Z. wrote the manuscript with input from other authors.

