## Supplementary figures and table for "Mechanistic insights into Phage SSB-activated bacterial Retron-Eco8 immunity"

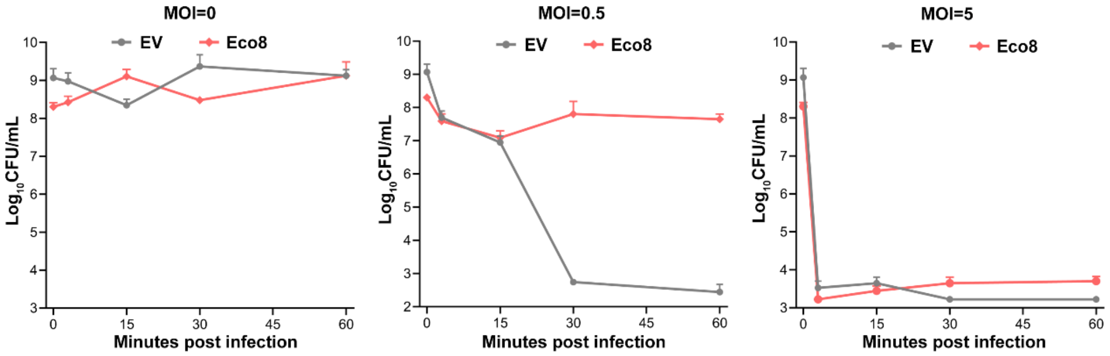


**Extended Data Fig. 1: Activation of the Retron-Eco8 system leads to irreversible cell death.**

The survival status of *E. coli* MG1655 transformed with either the pACYC-Retron-Eco8 system (Eco8) or the empty pACYC vector (EV) was assessed by plating and counting colony-forming units (CFUs) measured at 2, 15, 30, and 60 minutes post-infection (T4 phages) at multiplicities of infection (MOI) of 0, 0.5 ,5. Data are representative of three biological replicates. (n = 3; mean ± SD).

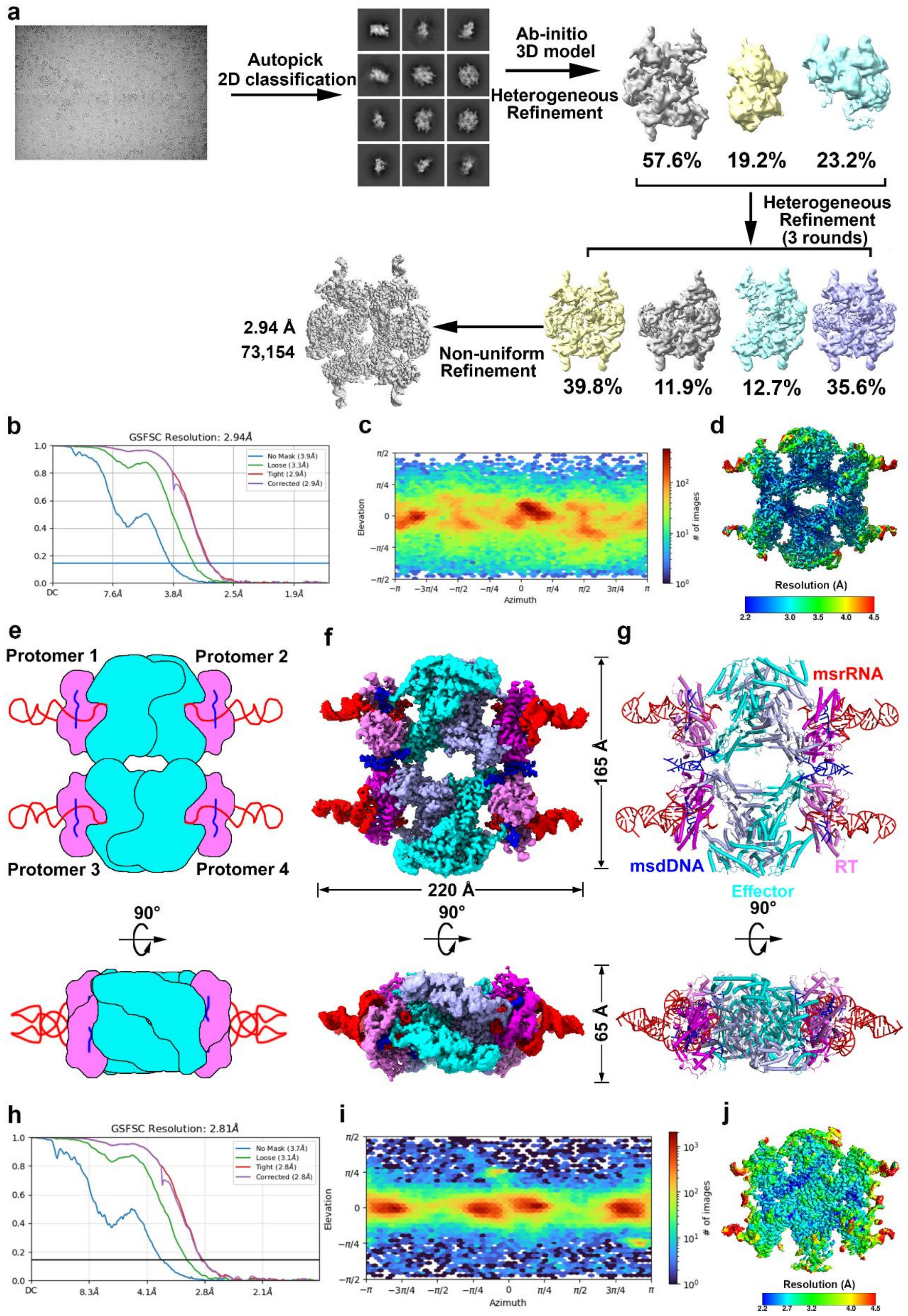


**Extended Data Fig. 2: Cryo-EM reconstruction of the Retron-Eco8 complex.**

**a** Flow chart of image processing for Retron-Eco8 complex.

**b, h** Fourier Shell Correlation curve of Retron-Eco8 (b) and Retron-Eco8$^{ATP}$ complexes (h).

**c, i** Direction distribution plot of Retron-Eco8 (c) and Retron-Eco8$^{ATP}$ complexes (i).
**d, j** Final 3D reconstructed map of Retron-Eco8 (d) and Retron-Eco8$^{ATP}$ complexes (j), colored according to local resolution.
**e-g** Schematic (e), Surface (f) and ribbon (g) representations of the 2.94-Å cryo-EM structure of the Retron-Eco8 complex.

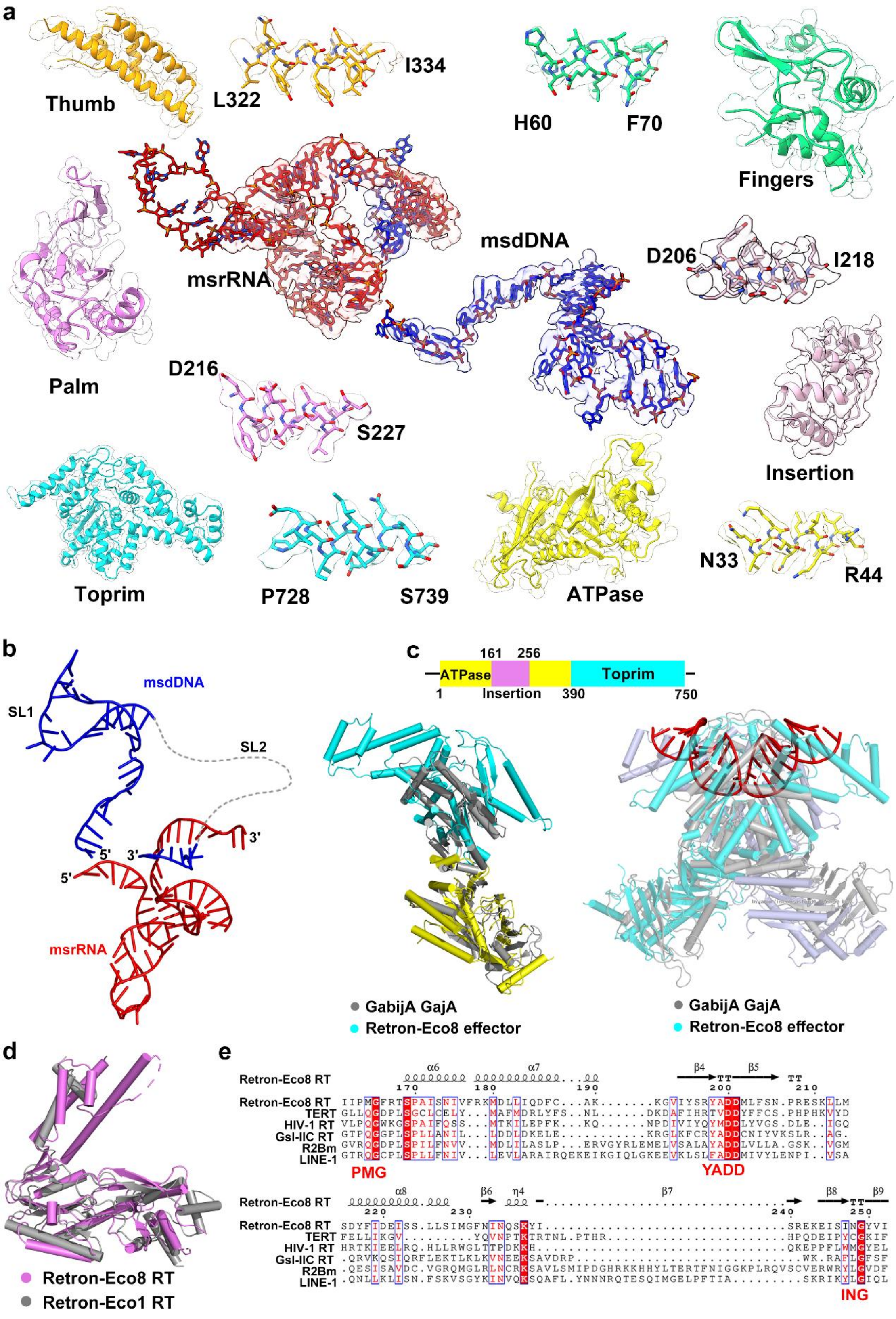


**Extended Data Fig. 3: Structural comparison of the effector and RT of Retron-Eco8 with related proteins.**

**a** Cryo-EM densities of each domain of Retron-Eco8. Densities for the indicated regions are shown in the context of the atomic model.

**b** Ribbon representation of the msrRNA-msdDNA for Retron-Eco8$^{ATP}$ complex.
**c** Schematic diagram showing the domain organization of effector protein within the Retron-Eco8 system (upper inset). Structural comparison of Retron-Eco8 effector (in color) with GajA (PDB, 8JQ9, in grey) (monomer, lower left; dimer, lower right).
**d** Structural comparison of Retron-Eco8 RT (in violet) with Retron-Eco1 RT (7V9U, in grey).
**e** Multiple sequence alignment of conserved catalytic residues in the RT of WP_053898075.1 (*Escherichia coli* 200499), NP_001035796(*Tribolium castaneum*), AAA44198.1(*Human immunodeficiency virus* 1), ADK35363.1(*Geobacillus stearothermophilus*), AAB59214(*Bombyx mori*), AAC51271.1(*Homo sapiens*). The conserved residues are indicated with red shading.

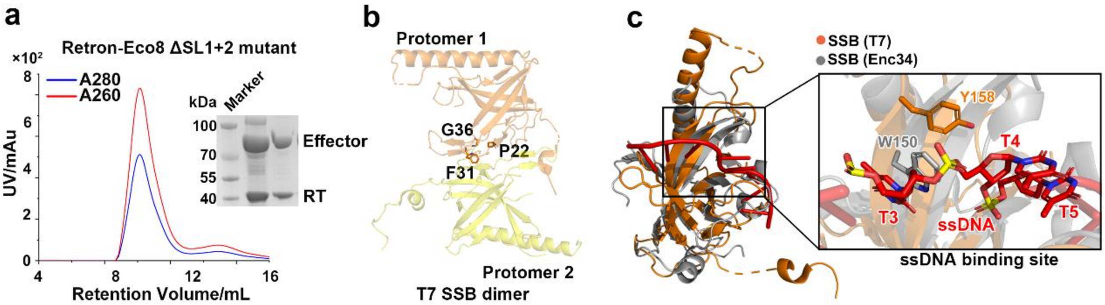


**Extended Data Fig. 4: T7 SSB recognizes the SLs region of the msdDNA.**

**a** SEC and SDS-PAGE profiles of Retron-Eco8 ΔSL1+2 mutant.

**b** Ribbon representation of the T7 SSB dimer (PDB, 1JE5). Three previously reported mutants which alter the oligomerization state are labeled.

**c** Overall and detailed structural comparison of T7 SSB (PDB, 1JE5) with phage Enc34 SSB (PDB, 5ODL).

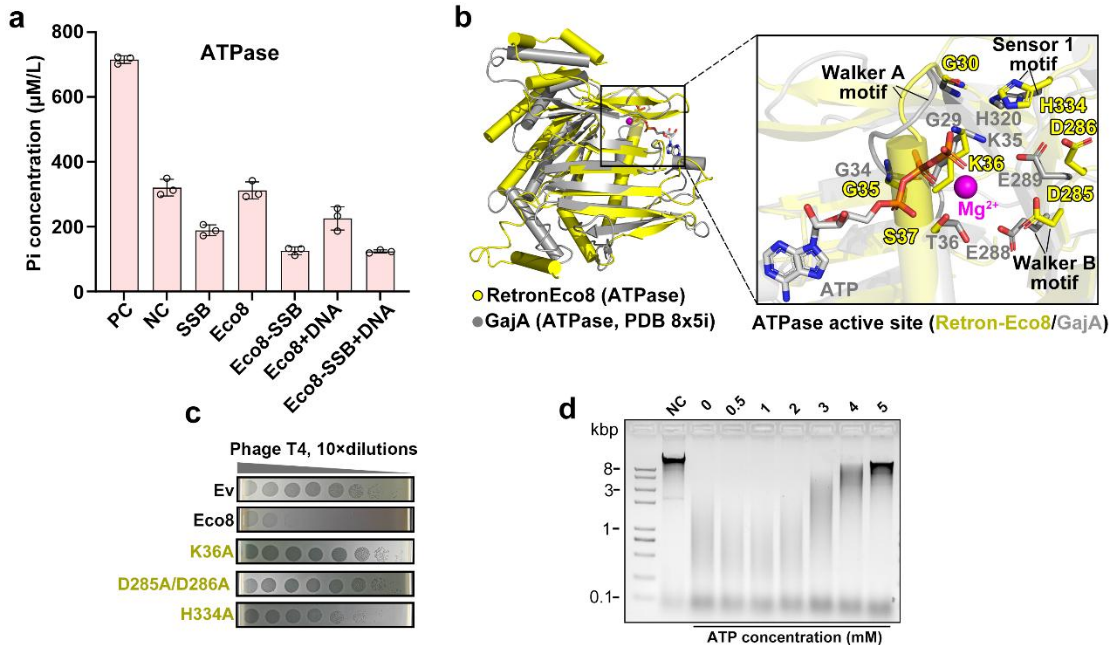


**Extended Data Fig. 5: The ATPase domain is critical for Retron-Eco8 to perform its function.**

**a** ATPase activity of the Retron-Eco8 system. Azaca system (*Bacillus massilioanorexius* AP8, IMG Genome ID: 2547132168) was used as a positive control. DNA: *E.coli* genomic DNA. Data are representative of three biological replicates. (n = 3; mean ± SD).

**b** Structural comparison of ATPase domain from Retron-Eco8 and GajA (PDB, 8X5I). Key residues are labeled.

**c** Serial dilution plaque assays for T4 phage on *E. coli* MG1655 strain transformed with plasmids encoding WT or indicated effector ATPase active site mutants.

**d** Nuclease activity assays (substrate: T4 genomic DNA) of Retron-Eco8-SSB complex in the presence of different concentration of ATP.

**Supplementary information Table S1. Cryo-EM data collection, refinement and validation statistics**

| | Retron-Eco8 | Retron-Eco8$^{ATP}$ | Retron-Eco8-SSB |
|---|---|---|---|
| **Data collection and processing** | | | |
| Magnification | 105,000 | 105,000 | 105,000 |
| Voltage (keV) | 300 | 300 | 300 |
| Electron exposure ($e^{-}$/Å$^2$) | 50 | 50 | 50 |
| Defocus range (μm) | 1.5 to 2.5 | 1.5 to 2.5 | 1.5 to 2.5 |
| Pixel size (Å) | 0.827 | 0.827 | 1.1 |
| Symmetry imposed | C1 | C1 | C1 |
| Initial particle images (no.) | 237,256 | 588,563 | 567,993 |
| Final particle images (no.) | 73,154 | 140,497 | 72,272 |
| Map resolution (Å) | 2.94 | 2.81 | 2.80 |
| FSC threshold | 0.143 | 0.143 | 0.143 |
| Map resolution range (Å) | 1.6-999 | 1.6-999 | 2.2-999 |
| **Refinement** | | | |
| Initial model used (PDB code) | *ab-initio* | *ab-initio* | |
| Model resolution (Å) | 2.69 | 3.25 | |
| FSC threshold | 0.5 | 0.5 | |
| Map sharpening *B* factor (Å$^2$) | -73.0 | -63.6 | -60.0 |
| Map Correlation Coefficient | 0.81 | 0.65 | |
| **Model composition** | | | |
| Non-hydrogen atoms | 36,872 | 41,900 | |
| Protein residues | 3,760 | 4,084 | |
| Nucleotides | 288 | 400 | |
| ***B* factor (Å$^2$)** | | | |
| Protein | 68.58 | 68.75 | |
| Nucleotides | 123.41 | 46.64 | |
| **R.m.s. deviations** | | | |
| Bond lengths (Å) | 0.003 | 0.003 | |
| Bond angles (°) | 0.583 | 0.643 | |
| **Validation** | | | |
| MolProbity score | 1.67 | 1.74 | |
| Clash score | 8.08 | 13.02 | |
| Poor rotamers (%) | 0.66 | 0.21 | |
| **Ramachandran plot** | | | |
| Favored (%) | 96.53 | 97.41 | |
| Allowed (%) | 3.47 | 2.59 | |
| Disallowed (%) | 0.00 | 0.00 | |
| **EMDB** | EMD-62954 | EMD-66110 | EMD-63117 |
| **PDB** | 9LBQ | 9WN8 | |
